# DNA-FISH Metaphase Spreads to Distinguish Extrachromosomal DNA from Homogeneously Staining Regions in Human Cancer Cell Lines

**DOI:** 10.64898/2026.07.07.735342

**Authors:** Lauren Marie Masters, Katelin Maria Hagstrom, Graham Scott Erwin

## Abstract

Whole-genome sequencing identifies focal DNA amplifications with base-pair resolution but cannot determine whether amplified sequences reside on extrachromosomal DNA (ecDNA, also known as double minutes) or within chromosomally integrated homogeneously staining regions (HSRs). DNA fluorescence in situ hybridization (DNA-FISH) metaphase spreads remain the gold standard for distinguishing these amplification states at single-cell resolution. Here, we present a detailed protocol for DNA-FISH metaphase spreads using human cancer cell lines, encompassing cell culture, metaphase arrest, hypotonic treatment, fixation, chromosome spreading, fluorescent probe hybridization, and fluorescence imaging. The protocol incorporates intermediate quality-control steps to verify successful chromosome dispersion and optimize metaphase spread quality, making the workflow accessible to laboratories without specialized cytogenetics expertise. Results demonstrate clear visualization of ecDNA and HSR amplification states using locus-specific probes and illustrate common technical artifacts that can affect interpretation. This protocol provides a robust and reproducible approach for studying the structural organization of oncogene amplification in cancer cells.

**SUMMARY:** We report a DNA-FISH metaphase spread protocol that visually detects locus copy number and location within the genome. This approach enables single-cell resolution of amplification states, specifically in cancer cell lines containing extrachromosomal DNA and homogeneously staining regions.

## INTRODUCTION

The human genome is organized not only by its sequence, but also by its structure, and changes in either sequence or structure can lead to cancer^1–4^. Oncogene amplification is present in approximately 40% of all tumors, where it contributes to tumor initiation, growth, metastasis, and drug resistance^5^. In some cases, amplified oncogenes are carried on extrachromosomal DNAs (ecDNAs, also called double minutes), which are circular, self-replicating, double-stranded DNA elements that reside outside of the linear chromosomes^6–9^. ecDNAs are hundreds of kilobases to megabases in size^10,11^. Due to their lack of centromeres, they segregate asymmetrically during mitosis, leading to as many as 100+ copies in a single nucleus^12–18^. This uneven segregation generates copy number variability across daughter cells, which drives rapid tumor evolution and poor patient outcome^17,19^.

ecDNAs exist in a dynamic relationship with homogeneously staining regions (HSRs) (**Figure 1A**)^8,18^. HSRs are focal, intrachromosomal amplifications that can arise through the reintegration of ecDNAs into a single locus in the genome^8,10,18,20–31^. These two states, ecDNAs and HSRs, can interconvert, revealing that oncogene amplification is structurally plastic^24,25,32,33^. This interconversion has therapeutic relevance because ecDNAs can transition to the HSR state in the presence of drugs^32–38^. Cells that undergo this transition to the HSR state then become resistant to the drug under therapeutic pressure^33^. HSRs also appear to be more frequent in malignant tumors than in benign tumors^39^. Despite their clinical relevance, no FDA-approved strategies target the HSR state, in part because methods to distinguish these amplification structures remain underutilized.

**Figure 1:**
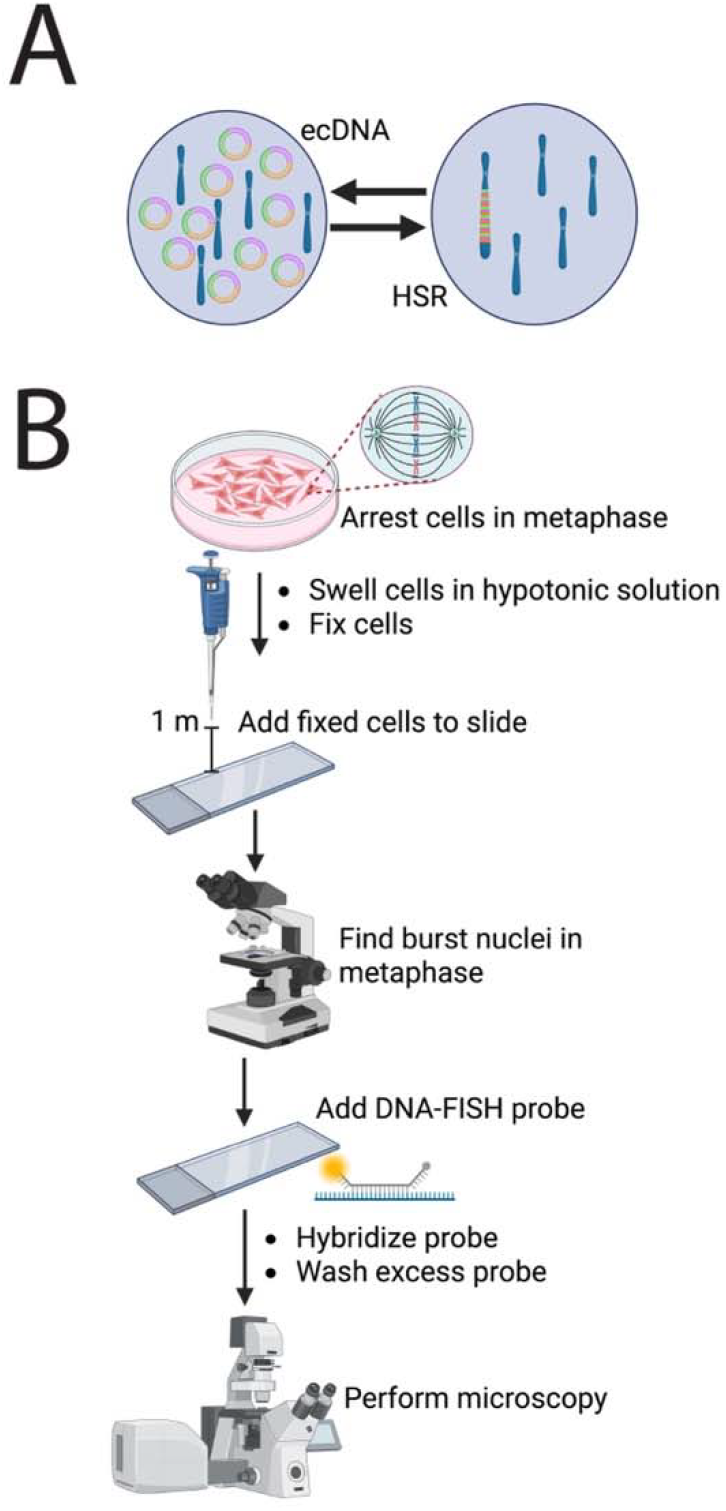
Overview of ecDNA and HSR amplification states and the DNA-FISH metaphase spread workflow. (**A**) Schematic illustrating the interconversion between extrachromosomal DNA (ecDNA), also known as double minutes, and homogeneously staining region (HSR) amplification states. (**B**) Workflow for DNA-FISH metaphase spreads. Cells are arrested in metaphase, swollen in a hypotonic solution, and fixed. Fixed cells are then dropped onto a glass slide from a height of 1 m. Upon impact, nuclei burst, and chromosomes disperse across the slide surface. Dispersed chromosomes are confirmed by phase-contrast microscopy. A DNA-FISH probe is then applied and hybridized to the target DNA locus. The hybridized signal is subsequently visualized by fluorescence microscopy.

Although ecDNA and HSR states have distinct clinical consequences, current sequencing approaches cannot reliably distinguish them. Whole-genome sequencing (WGS) now identifies amplified regions with base-pair resolution^40^. WGS has been valuable in defining the sequence composition of ecDNAs and HSRs, but at the expense of spatial resolution. Unlike traditional cytogenetic approaches, WGS alone cannot determine whether an amplified sequence resides on ecDNAs or is integrated into a chromosome as an HSR^41–43^. As a result, methods that preserve the spatial context of amplification remain essential for understanding the biology of these distinct states.

For the past 40 years, DNA fluorescence in situ hybridization (DNA-FISH) metaphase spreads have served as the gold standard for identifying and distinguishing ecDNA and HSR amplification states^42,44–46^. In fact, ecDNAs and HSRs were first identified and characterized through metaphase-based cytogenetic analyses^7,47^. Before the widespread use of fluorescent probes, in situ hybridization was performed using radiolabeled DNA probes, which have also been used to visualize ecDNAs and HSRs^48,49^. As WGS gained in popularity, DNA-FISH metaphase spreads became less common. Many aspects of the workflow are scattered across the cytogenetic literature, and optimized protocols are typically retained within individual research groups or cytogenetics labs that treat the method as proprietary^50,51^. Here, we describe a DNA-FISH metaphase spread protocol to visually resolve amplification state at the single-cell level.

## PROTOCOL

All human cell culture procedures were performed in compliance with institutional guidelines.

### 1. Arrest cells in metaphase using Colcemid (N-methyl-N-deacetyl-colchicine)

1.1. Begin with approximately 2.0 × 10□–2.0 × 10□ cancer cells or other cultured cells. Too few cells will yield fewer metaphase spreads, whereas too many cells may result in insufficient fixation.
1.2. Determine the appropriate Colcemid incubation time for the experimental system. Incubation time affects the number of metaphase spreads obtained and chromosome length, and optimal conditions vary among cell types. NOTE: Shorter incubation times result in fewer cells arrested in metaphase at the time of collection. Longer incubation times produce more condensed chromosomes with shorter arms, making analysis more difficult. For cancer cell lines with an approximate doubling time of 24 h, incubation periods ranging from 45 min to overnight produce suitable metaphase spreads. Optimize the incubation time for cells with slower doubling times. A 2 h incubation is generally sufficient to produce an adequate number of metaphase spreads with chromosome lengths suitable for analysis.
1.3. Calculate the amount of Colcemid required to achieve a final concentration of 10 μL/mL in the appropriate cell culture medium. NOTE: Maintain cells in logarithmic growth by passaging or seeding them at least 24 h before metaphase arrest. Treat cells with Colcemid when cultures reach 70%–90% confluence.
1.4. Dilute the Colcemid stock solution 1:1 with sterile ultrapure water. Add the appropriate volume to the culture and incubate for 45 min to overnight in a cell culture incubator.

### 2. Harvest cells

2.1. Prepare fresh Carnoy’s fixative (3:1 methanol: acetic acid). Keep the fixative on ice throughout the fixation procedure.
2.2. Prepare a fresh 0.075 M KCl solution and warm it in a 37 °C water bath.
2.3. Collect the cells as a pellet and record the cell count. Use this value to calculate the final resuspension volume. NOTE: Detach adherent cells by trypsinization. Transfer suspension cells directly into 15 mL centrifuge tubes. Dispose of Colcemid and Colcemid-containing media according to institutional hazardous waste guidelines.
2.4. Wash the cells with PBS.
2.5. Gently resuspend the cells in 10 mL of warm 0.075 M KCl and mix by gentle inversion. This hypotonic treatment swells the cells, allowing them to rupture and disperse chromosomes during slide preparation.
2.6. Incubate the cells in a 37 °C water bath for 30 min.
2.7. Slowly add 2 mL of ice-cold Carnoy’s fixative while gently mixing the suspension by inversion.
2.8. Allow the cells to stand for 5 min.
2.9. Centrifuge at 500 × *g* for 5 min at room temperature (RT).
2.10. Carefully aspirate the supernatant or remove it with a pipette.
2.11. Slowly add 10 mL of ice-cold Carnoy’s fixative in 0.5 mL increments while gently mixing after each addition.
2.12. Allow the cells to stand for 10 min at RT.
2.13. Centrifuge at 650 × *g* for 5 min at RT.
2.14. Repeat steps 2.10–2.13 twice
2.15. During the final wash, leave the cells suspended in Carnoy’s fixative.

PAUSE: Store the fixed cell suspension at −20 °C or proceed immediately to Section 3.

NOTE: For samples containing fewer than approximately 3 × 10L cells, perform fixation in 1.5 mL microcentrifuge tubes instead of larger conical tubes. Reduce the volumes of 0.075 M KCl and Carnoy’s fixative 10-fold while maintaining all incubation times, temperatures, and centrifugation conditions.

### 3. Tabletop dropping

3.1. Retrieve the sample from −20 °C and thaw it to RT.
3.2. Centrifuge at 800 × *g* for 7 min at RT.
3.3. Label the glass slides and place them on ice.
3.4. Using the cell count obtained before fixation, calculate the volume of Carnoy’s fixative required to resuspend the pellet at a final concentration of approximately 5 × 10□ cells/mL. After centrifugation, carefully remove the supernatant with a pipette, leaving the calculated volume of fixative above the pellet.
3.5. Resuspend the fixed cells in Carnoy’s fixative using a P1000 pipette.
3.6. Transfer approximately 300 μL of the fixed-cell suspension into a 1.5 mL microcentrifuge tube.
3.7. Thoroughly resuspend the sample using a P200 pipette by pipetting up and down several times.
3.8. Aspirate 60 μL of the resuspended cells using a P200 pipette.
3.9. Hold the pipette tip approximately 1 m above the glass slide on ice and aim for the center of the slide.
3.10. Slowly dispense one drop of the cell suspension onto the slide. NOTE: Mark the drop location on the edge of the slide with an ethanol-resistant marker to facilitate identification during subsequent steps.
3.11. Allow the cells to dry on ice for 10 min.
3.12. Place the slides on a slide incubator at 80 °C for 10 min.
3.13. Remove the slides with tongs and allow them to cool to the touch.
3.14. Examine the slides using a phase-contrast microscope with a 10× or 20× objective. Confirm the presence of multiple well-dispersed metaphase spreads without excessive overlap.

### 4. Pretreat the slide

4.1. Prepare Coplin jars for the wash steps
4.1.1. Prepare each reagent fresh: 70% acetic acid (40 mL), PBS (40 mL), PBS (45 mL), 70% ethanol (40 mL), 85% ethanol (40 mL), and 100% ethanol (40 mL).
4.2. Place the glass slides in 70% acetic acid for 2 min.
4.3. Place the glass slides in 40 mL of PBS for 30 s.
4.4. Place the glass slides in 45 mL of PBS for 5 min.
4.5. Dehydrate the glass slides sequentially in 70%, 85%, and 100% ethanol for 1 min each.
4.6. Air-dry the slides at RT for 10 min.

### 5. Prepare the probe

NOTE: This protocol uses commercially available locus-specific DNA-FISH probes that hybridize to a genomic region of interest.

5.1. Select a probe targeting the amplified locus to be examined. Thaw the probe and hybridization buffer according to the manufacturer’s instructions. Allow the probe to reach RT before use.
5.2. Prepare a probe mixture containing 1 μL of probe and 4 μL of hybridization buffer for each glass slide. NOTE: The optimal probe concentration may depend on the manufacturer and the probe. The concentration specified here has been optimized as per the probes used in this study. However, optimize the probe concentration for each new locus.
5.3. Apply 5 μL of the probe/hybridization buffer mixture to the center of each drop location on the glass slide. NOTE: Probes are light-sensitive. Minimize light exposure throughout the remaining procedure by storing the slides in a dark drawer between steps and while processing other slides.
5.4. Carefully place a coverslip over the sample.
5.5. Seal the edges of the coverslip with rubber cement.
5.6. Place the glass slides in a humid chamber.
5.7. Hybridize the probe according to the manufacturer’s protocol. Allow hybridization to proceed overnight (12–18 h).

### 6. Post-hybridization washes, counterstaining, mounting, and imaging

6.1. Prepare Coplin jars for the wash steps
6.1.1. Prepare each reagent fresh: 2× SSC (40 mL), 0.4× SSC (40 mL), 70% ethanol (40 mL), 85% ethanol (40 mL), 100% ethanol (40 mL), and PBS (45 mL).
6.1.2. Prepare and store SSC as a 20× stock solution (3 M NaCl, 0.3 M sodium citrate, pH 7.0). To prepare 1 L, dissolve 175.3 g NaCl and 88.2 g sodium citrate in 1 L of ultrapure water. Dilute the stock to prepare 2× SSC and 0.4× SSC working solutions as needed.
6.2. Warm the 0.4× SSC solution to 72 °C in a water bath.
6.3. Remove the glass slides from the humid chamber and carefully remove the rubber cement.
6.4. Place the glass slides in 2× SSC for 2 min at RT. Carefully remove the coverslips.
6.5. Place the glass slides in prewarmed 0.4× SSC at 72 °C for 2 min.
6.6. Place the glass slides in 2× SSC for 1 min at RT.
6.7. Dehydrate the glass slides sequentially in 70%, 85%, and 100% ethanol for 1 min each.
6.8. Air-dry the slides at RT for 10 min in the dark.
6.9. Dilute 1 mg/mL DAPI 1:1000 in PBS to a final volume of 1 mL. NOTE: This aliquot can be stored at 4 °C for up to 1 week.
6.10. Pipette 10 μL of the diluted DAPI onto each glass slide at the marked drop location.
6.11. Carefully place a coverslip over the sample to ensure the DAPI solution spreads evenly.
6.12. Incubate the slides in the dark for 5–10 min.
6.13. Place the glass slides in PBS for 30 s.
6.14. Remove the coverslips.
6.15. Place the glass slides in PBS for 2 min.
6.16. Dehydrate the glass slides by briefly dipping them in 70%, 85%, and 100% ethanol for approximately 10 s each.
6.17. Air-dry the slides at RT for 10 min in the dark.
6.18. Add 15 μL of mounting medium to each glass slide, then place a coverslip on top.
6.19. Allow the mounting medium to spread evenly, then gently blot away any excess.
6.20. Seal the edges of the coverslip with nail polish.
6.21. Allow the nail polish to dry for 15 min at RT in the dark. PAUSE: Store the slides at 4 °C for long-term storage.
6.22. Image the slides. NOTE: Images were acquired using an inverted fluorescence microscope with a 60× oil-immersion objective.
6.23. Collect multiple z-stacks for each fluorescence channel to capture signal throughout the full depth of the metaphase spread.
6.24. Deconvolve the image stacks using a maximum-likelihood estimation algorithm to enhance signal clarity and reduce out-of-focus fluorescence.
6.25. Process and analyze the final images in ImageJ, and select the optimal focal plane for each metaphase spread before downstream analysis.

## REPRESENTATIVE RESULTS

This protocol (**Figure 1B**) yields clean, well-dispersed, intact metaphase chromosomes suitable for cytogenetic analysis, with chromosomes and DNA-FISH signals clearly visible by fluorescence microscopy. Using a brightfield/phase-contrast microscope at low magnification (10× or 20×) is an important quality-control step to verify that nuclei have burst and chromosomes have dispersed (**Figure 2**). Successful fixation and tabletop dropping should yield well-dispersed metaphase spreads with multiple visible chromosome bursts across the slide. The yield of high-quality metaphase spreads is cell line-dependent and is also influenced by the duration of Colcemid treatment. As a general benchmark, successful preparations typically yield approximately 3–5 cells that burst at metaphase per field of view using a 20× objective^52^. Suboptimal metaphase spreads often exhibit cell clumping, a low proportion of burst nuclei, and/or poor chromosome dispersion, all of which interfere with downstream interpretation.

**Figure 2:**
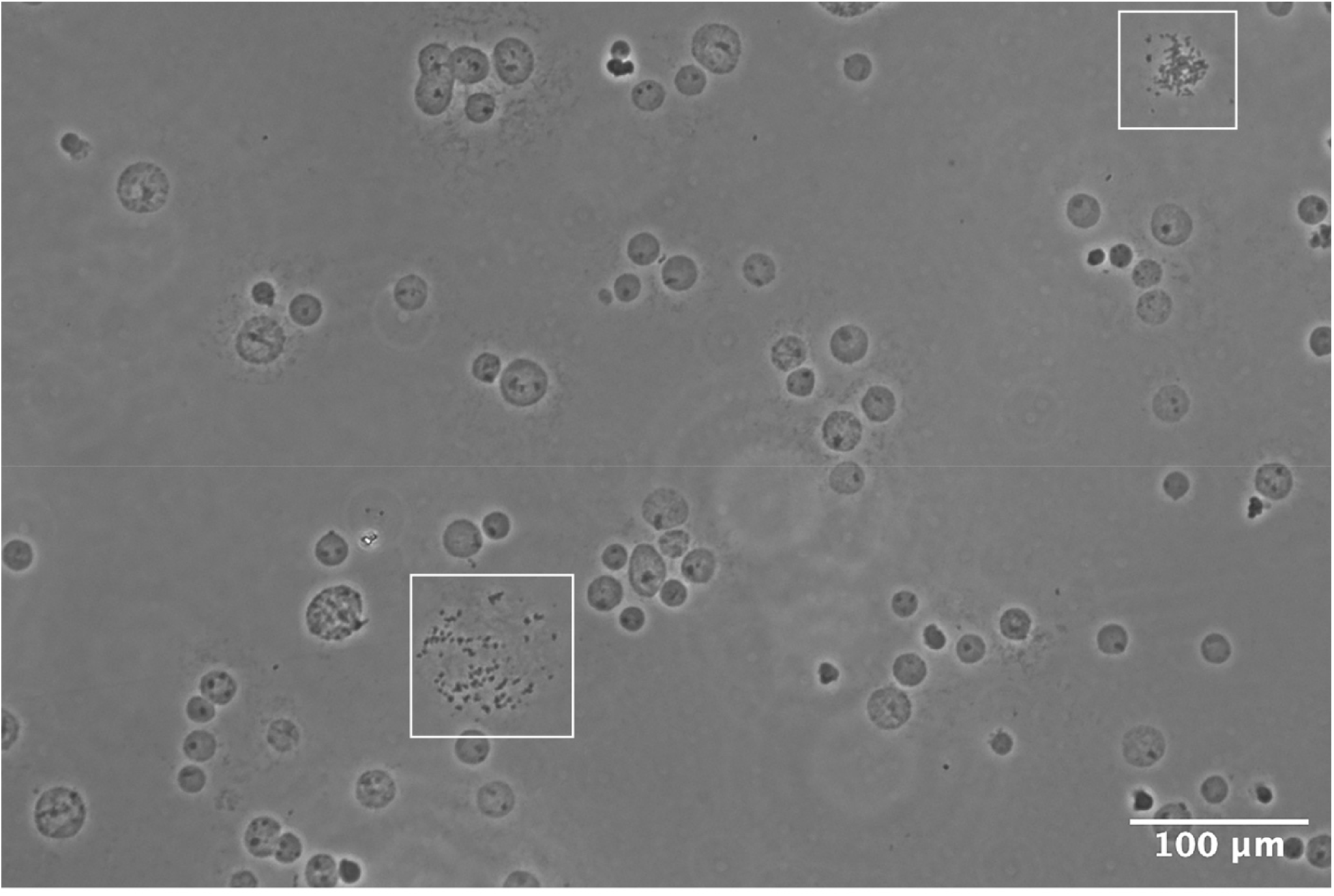
Confirmation of metaphase spreads by phase-contrast microscopy at 10× magnification. White boxes indicate burst nuclei with well-dispersed chromosomes, representing successful metaphase spread preparations. The presence of multiple burst nuclei across the field of view confirms successful metaphase arrest, hypotonic swelling, and nuclear rupture upon contact with the glass slide. Scale bar = 100 μm.

Representative DNA-FISH metaphase spread images illustrating ecDNA and HSR amplification states are shown in **Figure 3**. In the ecDNA state, the locus-specific probe should be detected outside the linear chromosomes (**Figure 3A,C**). Within a single sample, the number of ecDNA signals may vary substantially from cell to cell, reflecting the unequal distribution of ecDNA during cell division^12,13^. This heterogeneity is a characteristic feature of ecDNA-containing cell lines and should be expected.

**Figure 3:**
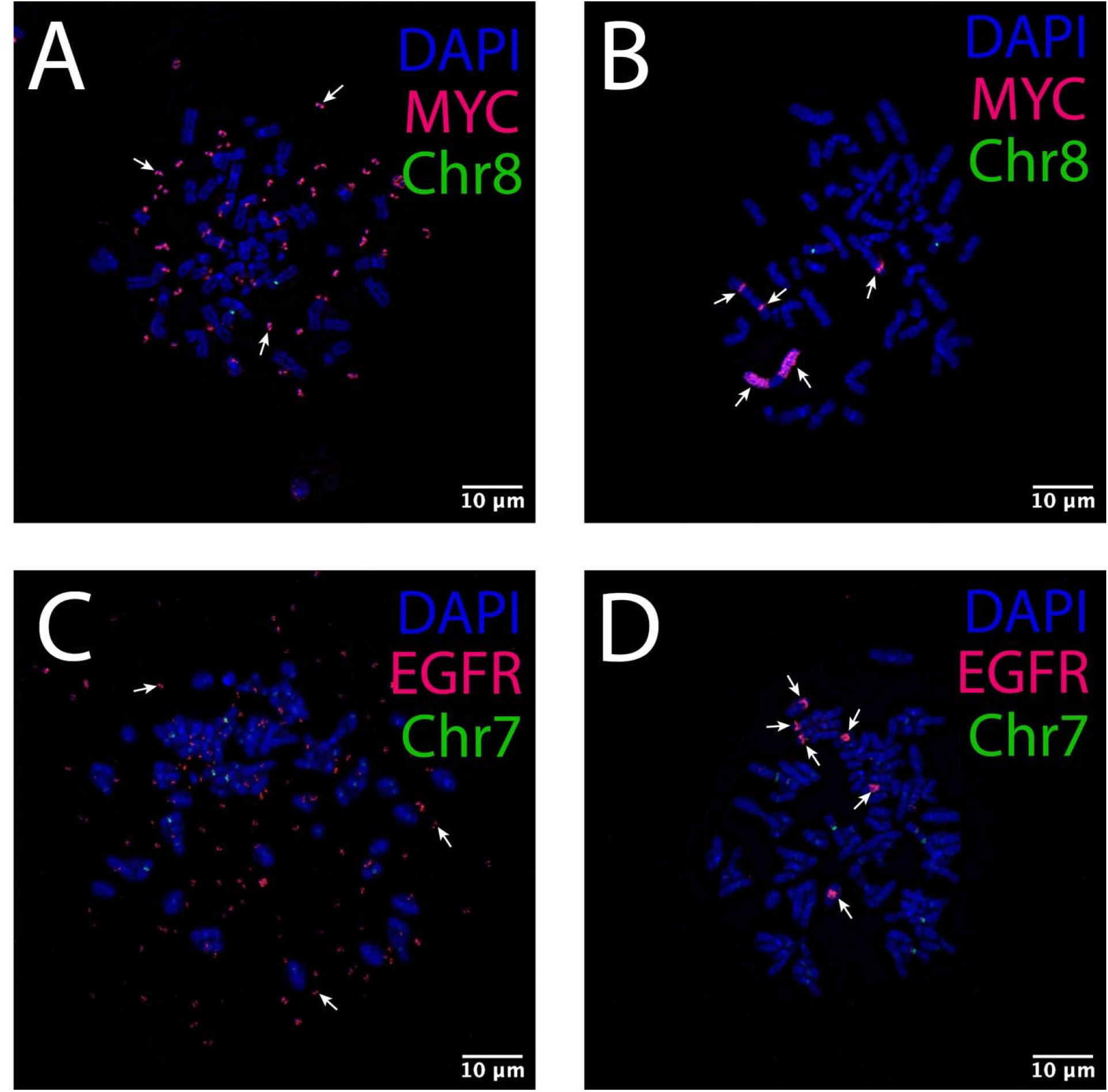
Representative DNA-FISH metaphase spread images distinguishing ecDNA and HSR amplification states. (**A**) DNA-FISH metaphase spread of a COLO320DM cell showing MYC signal (red) localized to ecDNA (examples indicated by white arrows); a chromosome 8 centromere probe (green) identifies the endogenous chromosome 8, where the native MYC locus resides. (**B**) DNA-FISH metaphase spread of a COLO320HSR cell showing MYC signal (red) localized to HSRs (examples indicated by white arrows); a chromosome 8 centromere probe (green) identifies the endogenous chromosome 8, where the native MYC locus resides. (**C**) DNA-FISH metaphase spread of a GBM39ec cell showing EGFR signal (red) localized to ecDNA (examples indicated by white arrows); a chromosome 7 centromere probe (green) identifies the endogenous chromosome 7, where the native EGFR locus resides. (**D**) DNA-FISH metaphase spread of a GBM39hsr cell showing EGFR signal (red) localized to HSRs (examples indicated by white arrows); a chromosome 7 centromere probe (green) identifies the endogenous chromosome 7, where the native EGFR locus resides. DAPI (blue) stains DNA in all panels. Scale bars = 10 μm.

In the HSR state, the probe signal appears as a dense cluster on a linear chromosome (**Figure 3B,D**). Unlike ecDNA, HSR signals are relatively consistent across nuclei within a single sample, reflecting the mitotic stability of chromosomally integrated amplification^3,13,18,53^. Because HSRs most often do not reside on the chromosome from which they originated, including a centromere probe for the endogenous chromosome is helpful. This centromere probe enables identification of both the native copy of the locus and the HSR-associated amplification. In some cell lines, multiple HSRs may be observed within the same metaphase spread, as demonstrated in GBM39hsr cells (**Figure 3D**). In addition, HSRs can vary considerably in size, with some appearing as small chromosomally integrated signals that may be difficult to distinguish from ecDNA without careful evaluation of their chromosomal localization (**Supplementary Figure 1**). Because cancer genomes vary substantially among tumor types and cell lines, ecDNA and HSR signal patterns may differ in abundance, chromosomal context, and overall appearance.

Representative images of common technical errors are shown in **Figure 4** to aid in the interpretation of unsuccessful preparations. In some cases, DNA condenses, but the nuclei fail to burst, making the resulting image difficult to interpret (**Figure 4A**). Nuclei that fail to burst are commonly observed, particularly when many cells have not entered metaphase. However, if fewer than 5–10% of nuclei burst, increasing the Colcemid incubation time, the drop height, or the KCl incubation may increase the number of successful metaphase spreads. In other cases, nuclei burst but do not complete metaphase arrest, resulting in uncondensed DNA that spills out of the cell and prevents accurate localization of the probe signal (**Figure 4B**). Increasing the Colcemid incubation time can increase the proportion of cells arrested in metaphase. However, excessive Colcemid exposure can lead to over condensed chromosomes with shortened chromosome arms^54^. Therefore, optimization of the Colcemid incubation time is essential. Poor chromosome spreading can also result in overlapping chromosomes, obscuring interpretation and making it difficult to determine the true location of the probe signal (**Figure 4C**). Reducing cell concentration in Carnoy’s fixative before tabletop dropping can decrease chromosome overlap.

**Figure 4:**
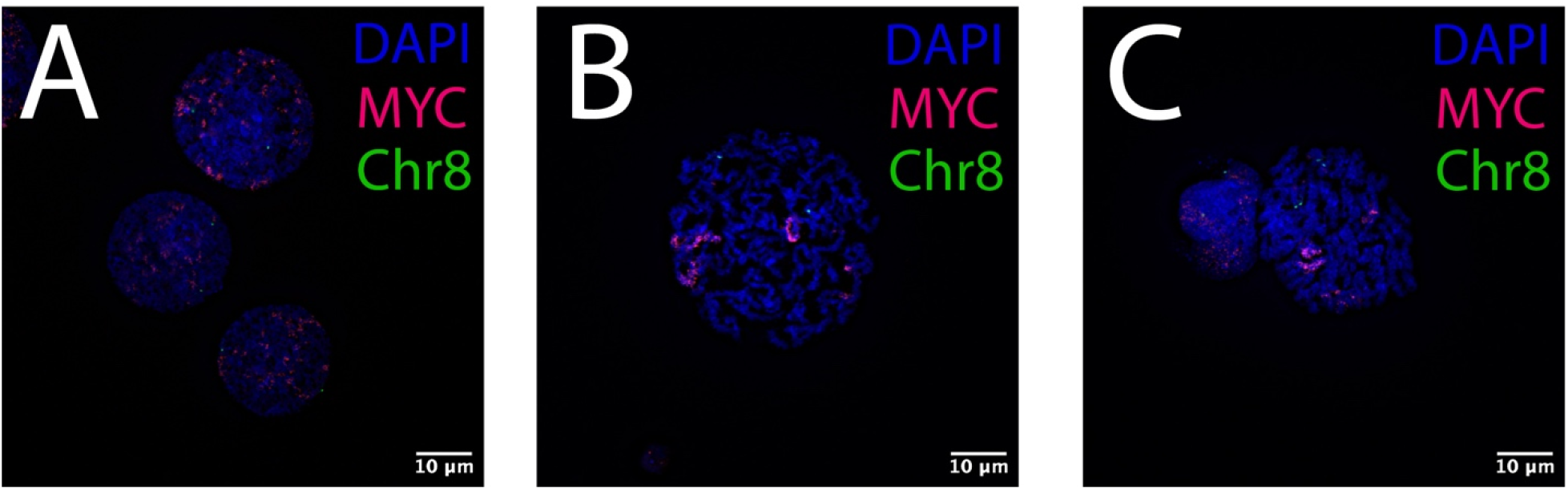
Representative images of unsuccessful metaphase spread preparations. (**A**) Example of COLO320DM nuclei that failed to burst, with DNA retained within intact nuclei. (**B**) Example of a COLO320HSR cell that failed to complete metaphase arrest, resulting in non-condensed DNA. (**C**) Example of two COLO320HSR nuclei that burst in close proximity, producing overlapping chromosomes. DAPI (blue), MYC (red), and chromosome 8 centromere probe (green). Scale bars = 10 μm.

When evaluating whether a signal represents true chromosomal integration, several criteria can help confirm the amplification state. First, if the signal is integrated into an HSR, it should appear on both sister chromatids because integrated sequences are replicated along with the host chromosome. Second, the signal should localize to the same focal plane as the DAPI-stained chromosome. Signals detected in a different focal plane may instead represent ecDNA overlapping the chromosome rather than true HSR integration. Finally, spectral karyotyping can be used to determine whether the locus has been stably integrated into the same chromosome across the cell population^55^.

Numerous approaches to metaphase spread preparation have been described^56–60^, and the method presented here represents one of many effective protocols. Important variables, including slide temperature, humidity, drop height, and cell concentration in the fixative, are sensitive to environmental conditions. To maintain consistent humidity within a laboratory, a cytogenetic drying chamber can be used during slide preparation. Individual optimization of these variables may be necessary to achieve consistent, high-quality metaphase spreads.

## DISCUSSION

Despite their well-established advantages, DNA-FISH metaphase spreads are underused in the era of WGS. The physical organization of amplified DNA is functionally significant, as it can influence downstream biological behavior and therapeutic response^61–64^. Focally amplified oncogenes can exist on ecDNA or within HSRs, and these two amplification states are not functionally equivalent^14,65–67^. For example, therapies targeting genes amplified on ecDNA can induce cell death, whereas cells that transition to an HSR state may remain viable and display drug resistance^33,34,39^. Accordingly, researchers need methods that reliably distinguish ecDNA from HSR amplification to understand how gene amplification contributes to cancer progression and treatment response.

In this approach, cells were first arrested in metaphase, generating highly condensed chromatin, allowing visualization of chromosome structures suitable for cytogenetic analysis^7,68^. To ensure chromosomes dispersed upon contact with the glass slide, cells were then swollen with a hypotonic solution. After multiple rounds of Carnoy’s fixation, cells were dropped onto slides from a height of one meter to burst nuclei and create metaphase chromosome spreads. Slides were then baked, washed, and hybridized overnight with a fluorescent DNA probe targeting a gene of interest, and counterstained with DAPI to visualize total DNA. Including a centromere probe for the endogenous chromosome containing the locus of interest helped identify the endogenous copy of the locus when present. This approach enabled the determination of whether a genomic locus was present on ecDNA, integrated within a chromosomal HSR, or present in other copy-number configurations.

Motivated by a longstanding interest in genome structure and repetitive DNA^69–72^, we sought to establish a robust DNA-FISH metaphase spread workflow for resolving amplification state. A major strength of this protocol is its emphasis on ensuring that nuclei burst effectively, producing well-dispersed chromosome spreads. For example, we were able to clearly visualize small integrated HSRs within COLO320DM cells (**Supplementary Figure 1**). These small HSRs have been previously reported^73,74^, but are easily overlooked when metaphase spreads are poorly dispersed or unclear. This protocol also describes intermediate validation steps during metaphase preparation to increase the likelihood of a successful outcome. Historically, expertise in metaphase preparation was transmitted by apprenticeship in cytogenetics laboratories. As research labs have focused more on sequencing and less on cytogenetics, fewer scientists possess this hands-on knowledge. This protocol aims to make the workflow more accessible to researchers without access to a cytogenetics lab.

While this technique distinguishes gene amplification states at the single-cell level, it has some limitations. DNA-FISH metaphase spreads do not provide exact copy number quantification for HSRs, as signal intensity reflects probe hybridization events rather than precise copy number. Additionally, prior knowledge of the locus of interest is required to select an appropriate probe. As a result, only the targeted locus will be visualized, and other potentially amplified regions will go undetected. DNA-FISH metaphase spreads also lack the base-pair resolution that sequencing provides. Finally, the protocol requires actively dividing cells that can be either maintained in culture or plated and incubated long enough to complete Colcemid treatment, depending on the cell type’s doubling time. Within these constraints, the protocol is compatible with a wide range of biological systems, including established cancer cell lines, primary tumor samples, and patient-derived cells, though optimization may be required for different cell types and model systems.

This protocol lowers the barrier to performing these analyses outside of a specialized cytogenetics laboratory. As more laboratories adopt DNA-FISH metaphase spreads, the field will be better positioned to determine when and why amplified oncogenes switch between ecDNA and HSR states.

## FIGURE AND TABLE LEGENDS

**Supplementary Figure 1:**
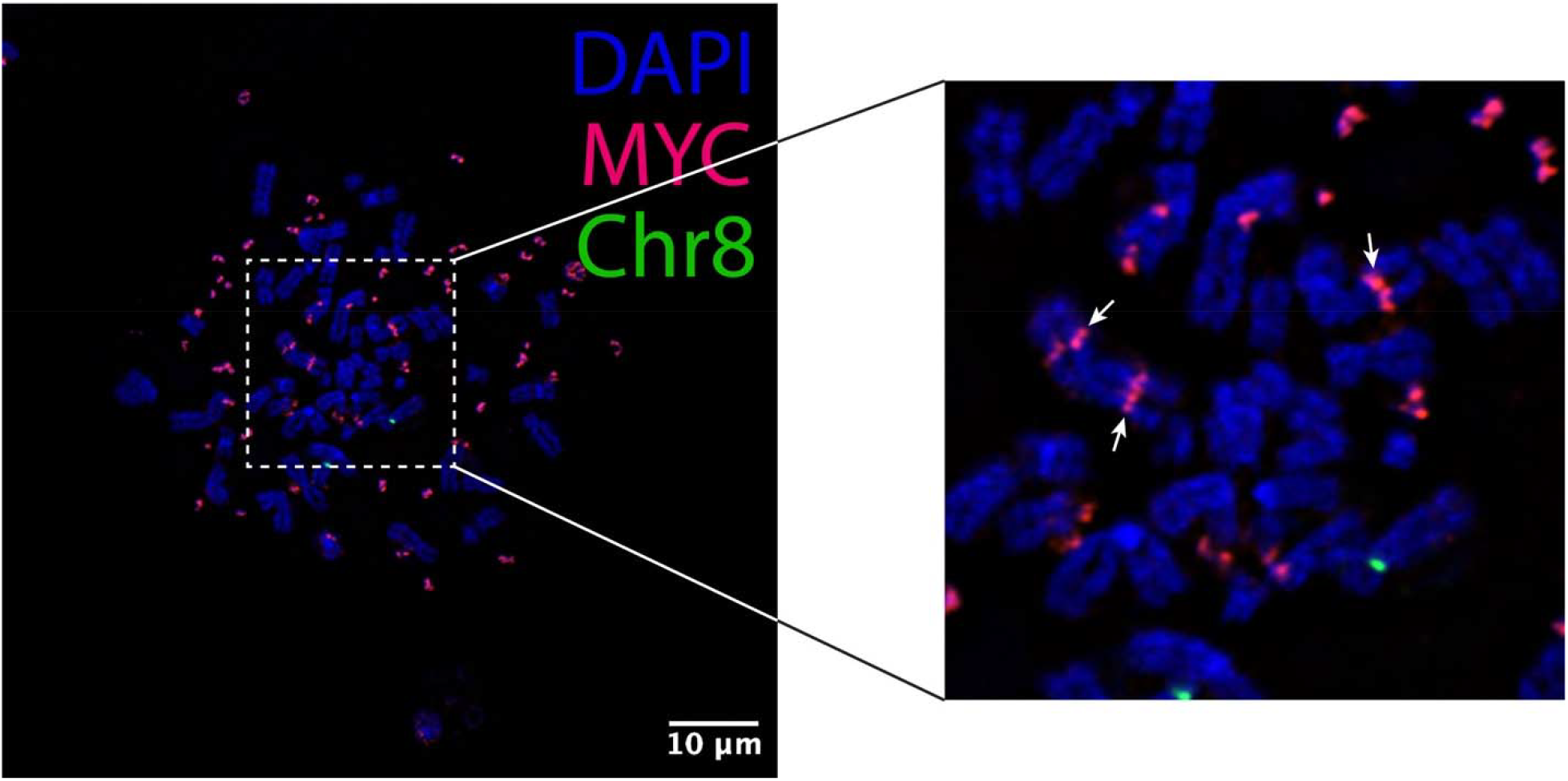
Small HSRs within the COLO320DM cell line. The dashed white box (left) indicates the region magnified on the right. White arrows indicate small HSRs integrated into linear chromosomes, which are readily visualized in well-dispersed metaphase spreads. Scale bars = 10 μm.

## ACKNOWLEDGMENTS

This work was supported by NIH R00HG011467 and CPRIT RR23004 to G.S.E. Imaging for this project was supported by the Integrated Microscopy Core at Baylor College of Medicine with funding from the NIH (P30 DK056338, P30 CA125123, S10OD030414). The authors thank Dr. Pawel Stankiewicz for his advice early on. The authors also thank Dr. Paul Mischel (Stanford University) for providing the GBM39ec and GBM39hsr cell lines.

