## Supplementary material for "DNA-FISH Metaphase Spreads to Distinguish Extrachromosomal DNA from Homogeneously Staining Regions in Human Cancer Cell Lines": Materials

| Name of Material/ Equipment | Company | Catalog Number | Comments/Description |
|---|---|---|---|
| 15-mL conical centrifuge | any source | | Centrifugation. |
| 15-mL conical tubes | any source | | Cell collection. |
| 200 proof ethanol | any source | | Wash steps. |
| Acetic Acid, glacial, 99.7% | Millipore Sigma | 695092 | Carnoys fixative and slide wash steps. |
| Antifade Mounting Medium | FisherScientific | NC9265087 | Slide mounting after completed slide prepation |
| Biological safety cabinet with vaccum outlet | any source | | Chemical storage. |
| Colcemid (*N* -methyl-*N* -deacetyl-colchicine) | Millipore Sigma | 10295892001 | Binds tubulin, arrests cells in metaphase. |
| Coplin jars | any source | | Wash steps. |
| DAPI (4′,6-diamidino-2-phenylindole) | Milipore Sigma | D8417-1MG | Fluorescent DNA stain. |
| EGFR/CON7 FISH probe | Empire Genomics | EGFR-CHR07 | Locus-specific flurescent probe. |
| Elmer's rubber cement | any source | | Keep coverslide in place during overnight probe hybridization. |
| Ethanol resistant permanent marker | any source | | Slide labeling. |
| fluorescent microscope | any source | | Slide imaging. |
| Glass Cover Slip, 22 x 22mm | ThermoFisher | 50-143-780 | Probe hybridization on slide. |
| Glass Slide, precleaned | VWR | 48311-703 | To drop metaphase spreads and perform DNA-FISH. |
| Humidified CO2 Incubator | any source | | Live-cell incubation. |
| Hybridization Buffer | Empire Genomics | Hyb-Buffer | Buffer used with probe. |
| ice bucket | any source | | Keep Carnoy's fixative cold. |
| KCl (2M) | ThermoFisher | AM9640G | Swell cells prior to fixation. |
| Methanol, 99.9%, HPLC grade | Millipore Sigma | 1060351000 | For Carnoys fixative. |

| | | | |
|---|---|---|---|
| microcentrifuge | any source | | Centrifugation. |
| microcentrifuge tubes | any source | | For cells during fixation, and for probe preparation. |
| MYC / SE 8 Probe | Leica | KI-10106 | Locus-specific flurescent probe. |
| PBS, pH 7.4 | Gibco | 10010-023 | Wash steps. |
| phase contrast microscope | any source | | Imaging of metaphase spreads immediately after dropping |
| Pipette tips | any source | | Pipetting. |
| sodium chloride | Millipore Sigma | S9888 | For SSC buffer. |
| sodium citrate | Millipore Sigma | 567446 | For SSC buffer. |
| sterile ultrapure water | any source | | Wash steps. |
| ThermoBrite programmable slide incubator | any source | | Slide incubation. |
| tongs | any source | | For slide wash steps. |
| water bath | any source | | For KCL incubation. |
